# Transcriptome analysis of *atad3-*null zebrafish embryos elucidates possible disease mechanisms

**DOI:** 10.1101/2024.11.02.621207

**Authors:** Shlomit Ezer, Nathan Ronin, Shira Yanovsky-Dagan, Shahar Rotem-Bamberger, Orli Halstuk, Yair Wexler, Zohar Ben-Moshe, Inbar Plaschkes, Hadar Benyamini, Ann Saada, Adi Inbal, Tamar Harel

**Author notes:** Corresponding author: Tamar Harel, M.D., Ph.D. Department of Genetics, Hadassah-Hebrew University Medical Center POB 12000, Jerusalem, Israel 9112001.

## Abstract

*ATAD3A*, a nuclear gene encoding the ATAD3A protein, has diverse roles in mitochondrial processes, encompassing mitochondrial dynamics, mitochondrial DNA maintenance, metabolic pathways and inter-organellar interactions. Pathogenic variants in this gene cause neurological diseases in humans with recognizable genotype-phenotype correlations. To further investigate the gene function and its implication in health and disease, we utilized CRISPR/Cas9 genome editing to generate a knockout (KO) model of the zebrafish ortholog gene, *atad3*. Analysis of *atad3*-null zebrafish embryos revealed microcephaly, small eyes, pericardial edema and musculature thinning, closely mirroring with the human rare disease phenotype. Larvae exhibited delayed hatching and embryonic lethality by 13 days post-fertilization (dpf). Locomotor activity, ATP content, mitochondrial content, and mitochondrial activity were all reduced in the mutant embryos. Transcriptome analysis at 3 dpf via RNA-sequencing indicated decline in most mitochondrial pathways, accompanied by a global upregulation of cytosolic tRNA synthetases, presumably secondary to mitochondrial stress and possibly endoplasmic reticulum (ER)-stress. Differential expression of select genes was corroborated in fibroblasts from an affected individual. The *atad3*-null zebrafish model emerges as a reliable representation of human *ATAD3A*-associated disorders, with similarities in differentially expressed pathways and processes. Furthermore, our study underscores mitochondrial dysfunction as the primary underlying pathogenic mechanism in *ATAD3A-*associated disorders and identifies potential readouts for therapeutic studies.

## 1. INTRODUCTION

*ATAD3A* is a nuclear gene that encodes ATPase family AAA domain-containing 3, a mitochondrial protein functioning as a hexamer spanning the inner and outer mitochondrial membranes. It is crucial to mitochondrial dynamics and influences the structure and function of the mitochondrial inner membrane and cristae (1), causing elongated or fragmented mitochondria when over- or under-expressed, respectively (2,3). Mice with a neuron-specific *Atad3* conditional knockout showed morphological changes in the mitochondria and cristae months prior to onset of neurological symptoms (1). Additionally, *ATAD3A* is implicated in mitochondrial DNA (mtDNA) maintenance (4). It binds Mitochondrial transcription factor A (TFAM), facilitating nucleoid trafficking along mitochondria and affecting respiratory complex formation (5). ATAD3A is also integral to cholesterol metabolism (6). Loss of *ATAD3A* results in accumulation of free cholesterol and triglycerides in hepatocytes, promoting nonalcoholic fatty liver disease (NAFLD) (7). In neurons, ATAD3A oligomerization induces cholesterol accumulation by inhibiting CYP46A1 (8). Furthermore, ATAD3A is involved in the interaction between endoplasmic reticulum (ER) and mitochondria (9,10), and is a crucial component in the mitochondria-associated membranes (MAM). It has a role in MAM integrity through interaction with the sigma-1 receptor, and disruptions can cause neurological pathogenesis (11). Aberrant ATAD3A oligomerization is also associated with poor MAM integrity (8). Elevated *ATAD3A* expression is correlated with poor prognosis in various cancers (12). In colorectal cancer, it provides chemoresistance by interacting with GRP78, suppressing ER-stress (13). In breast cancer, it prevents PD-L1 mitochondrial distribution through PINK1 inhibition, conferring tumor resistance (14). ATAD3A also regulates mitophagy by reducing PINK1 stability (15). Finally, ATAD3A has a role in the immune cGAS-STING dependent response, as pathogenic variants in this gene cause an upregulation of interferon-stimulated gene (ISG) expression (16).

This gene is one of three paralogs located in tandem on chromosome 1 in humans: *ATAD3A*, *ATAD3B* and *ATAD3C*. *ATAD3B* is expressed during embryonic development and in cancer cells and *ATAD3C* lacks four exons and is considered a pseudogene, leaving *ATAD3A* as the main, ubiquitously expressed gene out of the three (17). The high degree of similarity between the paralogs renders this genomic region excessively susceptible to non-allelic homologous recombination (NAHR) (3).

*ATAD3A* pathogenic variants cause distinct neurological diseases in humans with a predictable genotype-phenotype correlation. At one end of the spectrum are biallelic loss-of-function single nucleotide variants and biallelic deletions mediated by NAHR between the paralogs, which lead to a neonatal-lethal phenotype including severe brain malformations (i.e., simplified gyration patterns, white matter abnormalities and pontocerebellar hypoplasia) and respiratory failure (3,18,19). The reciprocal NAHR-mediated duplication gives rise to a similar lethal phenotype, associated with a dominant negative mechanism (20,21). At the other end of the spectrum is a recurrent dominant, monoallelic *de novo* variant (NM_001170535.3(*ATAD3A*): c.1582C>T, p.(Arg528Trp)) leading to developmental delay, optic atrophy, cardiomyopathy and peripheral neuropathy. A homologous mutation in the Drosophila *bor* gene, an ortholog to *ATAD3A* with about 70% similarity, demonstrated a similar phenotype acting in a dominant-negative manner (3). Another dominant-negative variant (NM_001170535.1(*ATAD3A*): c.1064G>A p.(Gly355Asp)) was described to lead to hereditary spastic paraplegia and axonal neuropathy (22). Elsewhere alongst the spectrum are biallelic hypomorphic variants in *ATAD3A,* whose clinical impact correlates with the severity of the mutation as modelled in the *Drosophila* gene *bor* (19).

Zebrafish are commonly used as models for many human diseases due to a relatively high degree of genetic resemblance between orthologs and their rapid development, which can be easily tracked in transparent embryos. Mitochondria generate up to 90% of cellular energy in the form of adenosine triphosphate (ATP) through oxidative phosphorylation; consequently, mitochondrial dysfunction is often apparent in tissues with high energy demand, such as the nervous system and muscles. Zebrafish offer an excellent model for studying mitochondrial diseases due to the well-studied development of their nervous system (23). Moreover, zebrafish models can be used for testing the phenotypic effect of different treatments relatively easily. The zebrafish ortholog gene, *atad3,* shares about 80% similarity to *ATAD3A*, and lacks paralogs. The protein domains of Atad3 are similar to both ATAD3A and ATAD3B. In order to further study the effects of *ATAD3A* variations, we used CRISPR/Cas9 genome editing technology to generate a zebrafish *atad3-*null model. We characterized the phenotype which was similar to the human disease and performed RNA sequencing (RNA-seq) to explore differentially expressed gene groups and pathways in the *atad3*-null zebrafish embryos. We describe reduced survival, downregulation of locomotor activity, and altered mitochondrial and metabolic pathways. Additionally, there was an upregulation of stress pathways including apoptosis and a global rise in cytosolic tRNA synthetases, which we propose is secondary to mitochondrial dysfunction and possibly ER-stress.

## 2. MATERIALS AND METHODS

### 2.1. Animal maintenance

We used AB/TL hybrid zebrafish (*Danio rerio*) as wild-type (WT). Adults were maintained according to standard procedures on a 14-h light/10-h dark cycle at 28°C. Embryos were produced by pair mating and raised at 28.5°C in egg water (0.3 gr NaCl in 1-liter distilled H_2_O). *atad3*^+/-^ zebrafish were kept on an AB/TL WT background. Guide for the Care and Use of Laboratory Animals guidelines were followed regarding animal maintenance and throughout all experiments presented in this work.

### 2.2. *atad3* expression in WT embryos

RNA was extracted from pools of WT embryos at different developmental stages (namely 6, 12, 24, and 48 hpf). cDNA was prepared using the qScript cDNA Synthesis Kit (Quantabio), and semi-quantitative reverse transcription polymerase chain reaction (RT-PCR) was performed on *atad3* compared to *eef1a1l1* as a housekeeping gene. Primers used are listed in Supplemental Table 1.

### 2.3. Whole-mount in situ hybridization

Whole-mount in situ hybridization using riboprobes for *atad3* (primers for sense and antisense probes listed in Supplemental Table 1) was performed according to standard protocol (Thisse and Thisse, 2008) on embryos at 48 hours post fertilization (hpf) using DIG RNA Labeling Kits (Roche, Basel, Switzerland). Anti-sense and sense RNA probes were synthesized using T7 or T3 RNA polymerases (Promega, USA). Sense probes served as negative control to each anti-sense probe.

### 2.4. Generation of *atad3*-KO in zebrafish by CRISPR/Cas9

CRISPR/Cas9 target sites were designed using CHOPCHOP tool (24) which identified crRNAs targeted to exons 1 and 2 of *atad3* (gd3 and gd1, respectively, listed in Supplemental Table 1). crRNAs were purchased from IDT. Each crRNA was injected separately to zebrafish eggs at the 1-cell stage according to IDT protocol to generate F0 mosaic embryos. To ensure efficacy, several F0 embryos were sacrificed and used for genomic DNA extraction and PCR amplification using primers spanning the CRISPR cleavage site and subsequent sequencing. The remaining F0 embryos were raised to adulthood. Fin clips were obtained at age 2 months for DNA extraction. DNA was amplified and sequenced to identify fish with a high degree of mosaicism, which were outcrossed to WT. Progeny of these crosses were genotyped to identify and characterize indels transmitted through the germline, and heterozygous F1 fish carrying desired mutations were outcrossed to generate the F2 generation.

### 2.5. Genotyping

DNA was extracted from fin clips of 2 months old zebrafish or from whole embryos as described (25).

DNA was amplified using primers spanning the CRISPR cleavage point. Genotyping for F1 and F2 generations was done by separation on a 15% acrylamide gel (Sigma) as previously described (26), and Sanger sequencing was used to correlate specific patterns on the acrylamide gel with the resulting indel (26). All primers used are listed in Supplemental Table 1.

### 2.6. Imaging and image analyses

Images of zebrafish embryos were acquired using Discovery.V8 stereoscope and AxioCam MRc digital camera (Zeiss) while embryos were mounted on their side in 0.5% low melting point agarose (Lonza, 50101) in 30% Danieau’s solution (1740 mM NaCl, 21 mM KCl, 12 mM MgSO4 · 7H2O, 18 mM Ca(NO3)2, 150 mM HEPES buffer) and 0.01% tricaine.

For phenotyping, embryos/larvae were kept individually over days 2-5 post fertilization (dpf), imaged daily and then genotyped as described. Circumference of head, eye and pericardial sac, length of neurocoel and tail thickness at somites 10 and 20 were measured using ImageJ software. Numbers vary throughout the days either due to death of embryos or, in some cases, since the angle in which embryos were mounted and imaged did not allow precise measurement.

### 2.7. Hatching assay

144 embryos were checked at different time points: 52hpf, 56hpf, and 72hpf. At each time point the hatched and unhatched embryos were separated to different plates, then tested for phenotype (mutants/siblings) and counted at 4 dpf.

### 2.8. Survival assay

89 embryos divided to three groups were checked daily starting at 3 dpf (when mutant phenotype begins to be clearly visible) until no mutant larvae survived. Each day the living mutants and siblings were counted, and dead larvae were taken out. Larvae were fed daily beginning at 5 dpf, and water was replaced about an hour after feeding.

### 2.9. Mitochondrial DNA content

Absolute mtDNA amount was quantified on extracted DNA by qPCR of the mitochondrial genes *mt-nd1, mt-nd6* and *mt-cox1* normalized to the nuclear gene *b2m*. DNA was pooled according to phenotype from three biological replicates, and a total of 10ng was used per reaction. Each mutant pool consisted of 3-8 mutants and each siblings’ pool consisted of 6-16 siblings. Quantification was performed using PerfeCTa SYBR Green FastMix (Quantabio). Three technical replicates were obtained for each sample. Primers are listed in Supplemental Table 1.

### 2.10. ATP content assay

Four biological replicates of 6 dpf zebrafish larvae were classified by phenotype and immersed in liquid nitrogen prior to sectioning of the heads. For each phenotype, 3 heads were collected and snap frozen in liquid nitrogen. Heads were kept in −80 degrees Celsius until assay. ATP level was quantified by the ATPlite® luciferin-luciferase bioluminescence assay according to the manufacturer’s instructions (Perkin Elmer Waltham MA, USA). Briefly, tissue was homogenized in Mammalian Cell Lysis solution and incubated with Substrate solution. Luminescence was read using Synergy HT BioTek microplate reader and relative luminescence units were normalized to protein. Total protein was quantified using Pierce™ Detergent Compatible Bradford Assay Reagent (ThermoFisher). Samples were collected on four different occasions, and the ATP content of each sample relative to the average ATP content in the samples from the occasion was calculated. Statistical analysis was carried out by a two-tailed, paired T-test.

### 2.11. Enzymatic assays

Three biological replicates of 5 dpf zebrafish larvae were classified by phenotype and immersed in liquid nitrogen prior to sectioning of the heads. For each phenotype, 7-9 heads were collected and frozen in −80 degrees Celsius until assay, then homogenates were prepared by sonication. The activities of respiratory chain complex IV (cytochrome *c* oxidase, COX), complex II+III and succinate dehydrogenase (SDH) were measured by spectrophotometry as previously described, using a double beam Kontron Uvikon 930 BioTek spectrophotometer (27,28). Enzymatic activities of COX and complex II+III were normalized to SDH activity and compared to the average activity of siblings. Statistical analysis was carried out by a two-tailed, paired T-test.

### 2.12. Locomotor activity assays

24 mutant larvae and 24 siblings were classified by phenotype at 4 dpf. At 5 dpf they were transferred to two 24-well plates, each of which contained 12 mutant larvae and 12 siblings. Each plate was then placed in a DanioVision Observation Chamber (Noldus), where the movement of the larvae was recorded for 6 hours and tracked using Noldus Ethovision XT16 (29). The experimental lighting conditions were as follows: 2 hours of white light, followed by 2 hours of darkness and 2 more hours of white light (2hrL:2hrD:2hrL). After 5 hours, the larvae were exposed to a tap to measure their acoustic startle response (30). The experiment was repeated with the same larvae the following day (at 6 dpf). The trajectory of each larva was smoothed by LOESS and segmented into periods of progressions and of stops (31). Responses to the tap and to turning off the light (light-off response, also known as visual motor response (32)) were measured by the distance moved (in cm) in the first 0.5 second after the stimulus.

Comparison of total activity (distance moved), of the tap response intensity and of the light-off response intensity between genotypes was performed using linear mixed models. In all models the genotype, the experimental day (5 dpf or 6 dpf) and the interaction between genotype and day served as fixed effects. Each larva was measured twice, and therefore the larval ID served along with the DanioVision chamber ID as random effects. Square root transformations were required for the models’ assumptions to hold. All post hoc analysis p-values obtained from all models were adjusted simultaneously for multiple comparisons using the BH procedure (33), controlling the false discovery rate at the 0.05 level. Statistical analysis was performed using a designated R code, utilizing the packages *“lmerTest”* (34) and *“emmeans”* (35).

### 2.13. RNA-seq

3 dpf (72hpf) embryos were separated into individual Eppendorf tubes and submerged in liquid nitrogen, and both RNA and DNA were extracted from each embryo using TRIzol Reagent (ThermoFisher) according to standard method, with 50µl TRIzol per embryo. DNA of single embryos was amplified and genotyped using acrylamide gel as described. Three biological replicates of RNA, each pooled from 5 mutants or WT embryos, were analyzed in technical triplicates by RNA-seq. Briefly, libraries were prepared using the KAPPA HyperPrep Plus RNA Library Prep Kit, and were then sequenced on a NovaSeq 6000 sequencing system (Illumina, San Diego, California, USA), generating 160bp paired-end reads.

### 2.14. Analysis of RNA-seq data

Raw reads were processed for quality trimming and adaptors removal using fastx_toolkit v0.0.14 and cutadapt v2.10 (36). The processed reads were aligned to the *Danio rerio* transcriptome and genome version GRCz11 with annotations from Ensembl release 99 using TopHat v2.1.1 (37). Counts per gene quantification was done with htseq-count v2.01 (38). Normalization and differential expression analysis were done with the DESeq2 package v 1.36.0 (39). Pair-wise comparisons were tested with default parameters (Wald test), without applying the independent filtering algorithm. Significance threshold was taken as padj<0.1. In addition, significant DE genes were further filtered by the log2FoldChange value. This filtering was baseMean-dependent and required a baseMean above 5 and an absolute log2FoldChange higher than 5/sqrt(baseMean) + 0.3 (for highly expressed genes this means a requirement for a fold-change of at least 1.2, while genes with a very low expression would need a 5.8-fold change to pass the filtering).

### 2.15. Gene Set Enrichment Analysis (GSEA)

Whole differential expression data were subjected to gene set enrichment analysis using GSEA (40). GSEA uses all differential expression data (cutoff independent) to determine whether a priori defined sets of genes show statistically significant, concordant differences between two biological states. We used the following gene sets collections: Gene Ontology Biological Process (GOBP), WikiPathways (WK) and Kyoto Encyclopedia of Genes and Genomes (KEGG), all downloaded from the FishEnrichr database (41).

### 2.16. RT-qPCR validations

For verification of differential expression on a group of selected genes, larvae at 5 dpf were individually genotyped. RNA of 9 homozygous WT larvae and 9 homozygous mutant larvae were pooled separately. For validation of RNA-seq data using RT-qPCR, 3 dpf embryos were divided to mutants (homozygous mutant) and siblings (phenotypically normal, representing homozygous WT and heterozygous embryos) and pooled. RNA was extracted from a pool of 15-20 embryos of each. cDNA was prepared using the qScript cDNA Synthesis Kit (Quantabio), and RT-qPCR was performed using PerfeCTa SYBR Green FastMix (Quantabio). The primers used for RNA differential expression validations are listed in Supplemental Table 1. Expression was normalized to *gapdh*.

### 2.17. Cell culture

Fibroblasts from an affected individual (reported in (3) and consented under research protocol H-29697) were maintained in DMEM Dulbeccòs Modified Eagle Media high glucose supplemented with 15% fetal bovine serum, 2mM L-glutamine and 1% penicillin-streptomycin (Biological Industries, Beit Haemek, ISRAEL).

### 2.18. Statistical Analysis

All RT-qPCR experiments were performed in triplicates, and in biological replicates as indicated in the specific method. For RT-qPCR and survival assay statistical analysis was carried out by two-tailed two-sample t-test. For ATP content and enzymatic activity statistical analysis was carried out by two-tailed, paired t-test, comparing biological replicates of mutants and siblings. For comparison of zebrafish embryo measurements (WT, heterozygotes, and mutants) statistical analysis was carried out by two-way ANOVA test. For hatching analysis, statistical analysis was carried out by chi-squared test. Statistical analysis for locomotor activity is as detailed above (section 2.12). *p* values < 0.05 were considered statistically significant for all tests.

## 3. RESULTS

### 3.1. Expression pattern of *atad3* in zebrafish embryos parallels tissues affected in human disease

*atad3*, the zebrafish ortholog, is highly similar to the human paralogs *ATAD3A* and *ATAD3B* in terms of protein sequence and domains. *ATAD3A* domains from Waters et al., 2023 (42) and Uniprot, as compared to *ATAD3B* and the zebrafish ortholog Atad3, are shown in Supplemental Fig.5. To evaluate whether zebrafish embryos would be a proper model for *ATAD3A*-associated disorders, we analyzed the expression levels of RNA extracted from whole embryos at various stages (6, 13, 26, and 48 hpf) by RT-PCR. *atad3* was increasingly expressed through the stages tested (Supplemental Fig. 1A). Following this, we set to determine by whole mount in situ hybridization the expression pattern in 2 dpf embryos. The strongest levels of detection were identified in specific regions of the brain and in the eyes (Supplemental Fig 1B). Since these organs/tissues are also those most affected by *ATAD3A*-associated disorders in humans, we pursued to generate knockout models in zebrafish through CRISPR/Cas9 genome editing.

### 3.2. CRISPR/Cas9-knockout of *atad3* leads to morphological defects

Two CRISPR/Cas9 lines were generated, designated gd1 (*huj25*) and gd3 (*huj24*). Gd1 fish carried a 14bp deletion and gd3 fish had a 4bp deletion (Fig. 1A, Supplemental Fig. 2A), in both cases causing a frameshift with a premature termination codon triggering nonsense mediated decay (NMD), as apparent by the 83% reduction (*p* < 0.001) in *atad3* mRNA (Fig. 1B). Mutant embryos could be easily recognized by 3 dpf, and became more distinct as they grew. Interestingly, the mutant phenotype closely correlates with the human phenotype described for *ATAD3A*-related disease. Notable phenotypic features included microcephaly, small eyes, pericardial edema, and thinning of the musculature in homozygous mutants as compared to siblings; for all measurements except for length, *p* < 0.001 from 3 dpf (Fig. 1C). The length of the neurocoel (extending from the head to the tail of the embryo) of the mutant and normal sibling embryos was not significantly different on day 2 nor 3, indicating that the small head and eyes of the mutant embryos were not part of an overall slower development. The swim bladder, which usually inflates starting at 4 dpf, failed to develop. WT and heterozygotes (combined referred to as “siblings” or sibs) were indistinguishable at any age (Fig. 1D-H). Measurements were performed on gd3 embryos. Gd1 showed a similar phenotype (Supplemental Fig. 2B).

**Figure 1.**
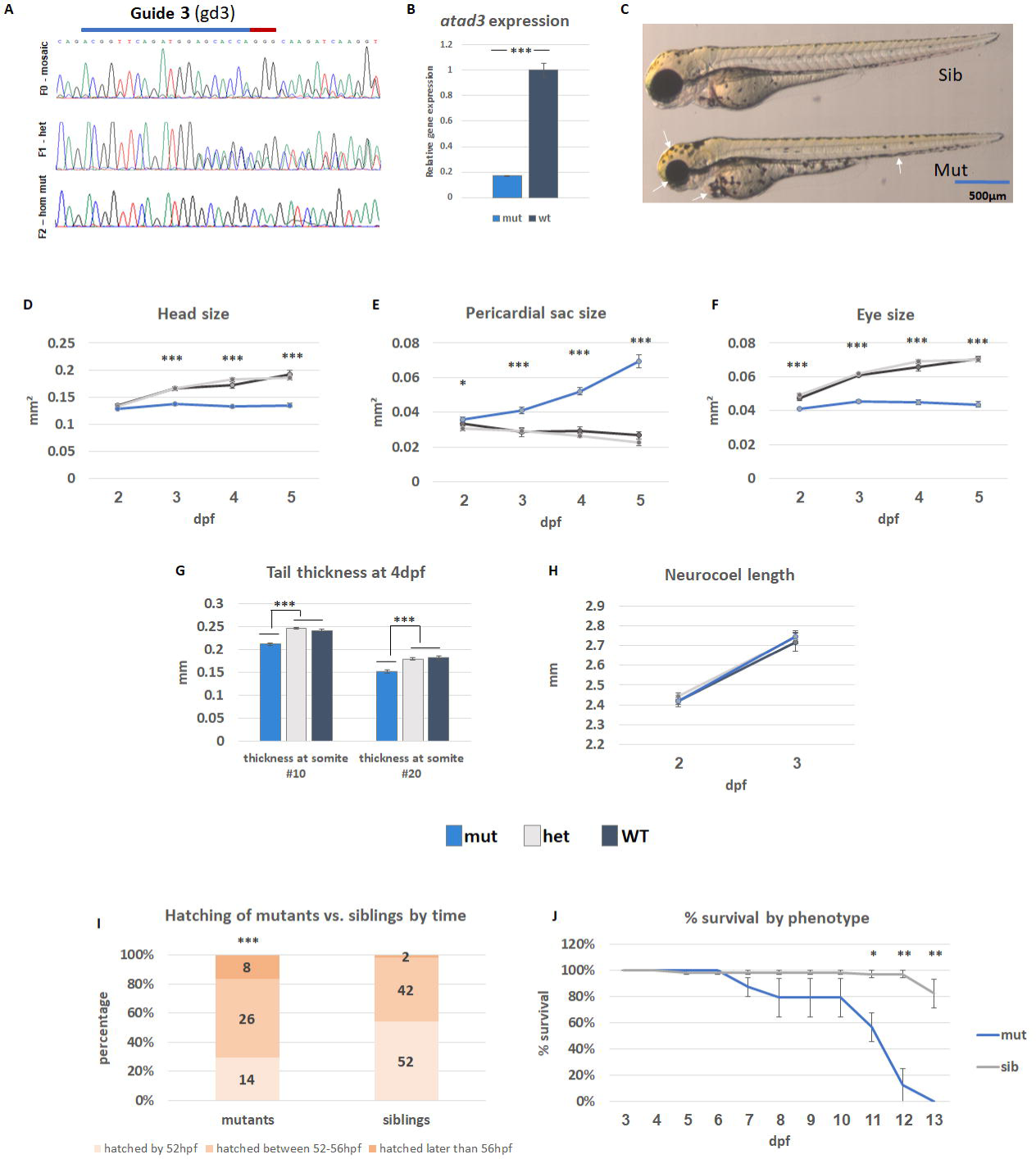
Characterization of *atad3*-null embryos/larvae. (A) DNA extracted from mutant embryos was sequenced to confirm a homozygous out-of-frame deletion. Upper panel – F0 generation, mosaic (CRISPR/Cas9-injected fish); middle panel – F1 generation (heterozygous); lower panel – homozygous mutant. CRISPR RNA (crRNA) marked by a blue line, PAM sequence marked by a red line. (B) *atad3* expression is drastically lower in RNA extracted from phenotypically mutant embryos compared to their siblings. (C-J) *Atad3*-KO embryos and larvae have a distinct phenotype. Embryos/larvae of two heterozygous gd3 fish were photographed throughout ages 2-5dpf and measured, then genotype was individually confirmed. (C) Siblings (either WT or heterozygous) vs. mutant embryo at 3dpf. The mutant embryos show smaller head and eyes, pericardial edema, thinning of the musculature and a similar overall length (indicated by white arrows). (D-H) measurements throughout ages 2-5dpf, error bars indicate standard error of means. Dark grey: WT; light grey: heterozygotes; blue: mutants. (D) Area of the head; € area of pericardial sac; (F) area of the eye; (G) tail thickness at 4dpf; (H) neurocoel length at 2 and 3 dpf. (I) Hatching time-range for 96 siblings and 48 mutants (*X*^2^ (1, *N* = 144) = 14.9, *p* =.000582, indicated above the mutant bar). Numbers on the bars indicate number of embryos in each group, Y axis shows the percentage of each group, for each phenotype. (J) Percent of larvae survival by day, according to phenotype: Mutants in blue (23 total, 7-8 in each biological replicate) and siblings in grey (66 total, 21-23 in each biological replicate). Error bars indicate the standard error of means of three biological replicates. Asterisks represent levels of significance (*p<0.05, **p<0.01, ***p<0.001).

### 3.3. *atad3*-null embryos hatch later than their siblings

The mutant embryos generally hatched later than the siblings, all within normal time range for zebrafish hatching. Among mutants, 29% hatched by the first time point (52hpf), 54% between the first and second (52-56hpf) and 17% after the second (>56hpf), while of the siblings, 54% hatched by the first time point, 44% between the first and the second and only 2% hatched after the third time point checked, showing a significant difference between mutants and siblings (*X*^2^ (1, *N* = 144) = 14.9, *p* =.000582) (Fig. 1I).

### 3.4. *atad3*-null larvae have reduced survival

Mutant larvae began to exhibit lethality at 7 dpf, and by 13 dpf none of the 23 mutant larvae survived. Conversely, survival rate in the sibling group (consisting of 66 embryos) was close to 100% until 12 dpf, with a statistically significant difference between the groups starting at 11 dpf (*p* = 0.022) and increasing throughout days 12 and 13 (*p* = 0.0027 and *p* = 0.0015, respectively) (Fig. 1J).

### 3.5. *atad3*-null larvae have decreased mitochondrial DNA content

Previously, pathogenic *atad3* variants in a *Drosophila* model were shown to have decreased mtDNA content, probably due to increased mitophagy (3). We therefore hypothesized that the overall mitochondrial content in zebrafish null larvae will be reduced. A mtDNA content assay, based on comparison of mitochondrial content normalized to nuclear content by qPCR, revealed a consistent and significant reduction of 19-23% (*p* < 0.001) in mtDNA content in mutant larvae compared to their phenotypically healthy siblings at 5 dpf (Fig. 2A).

**Figure 2.**
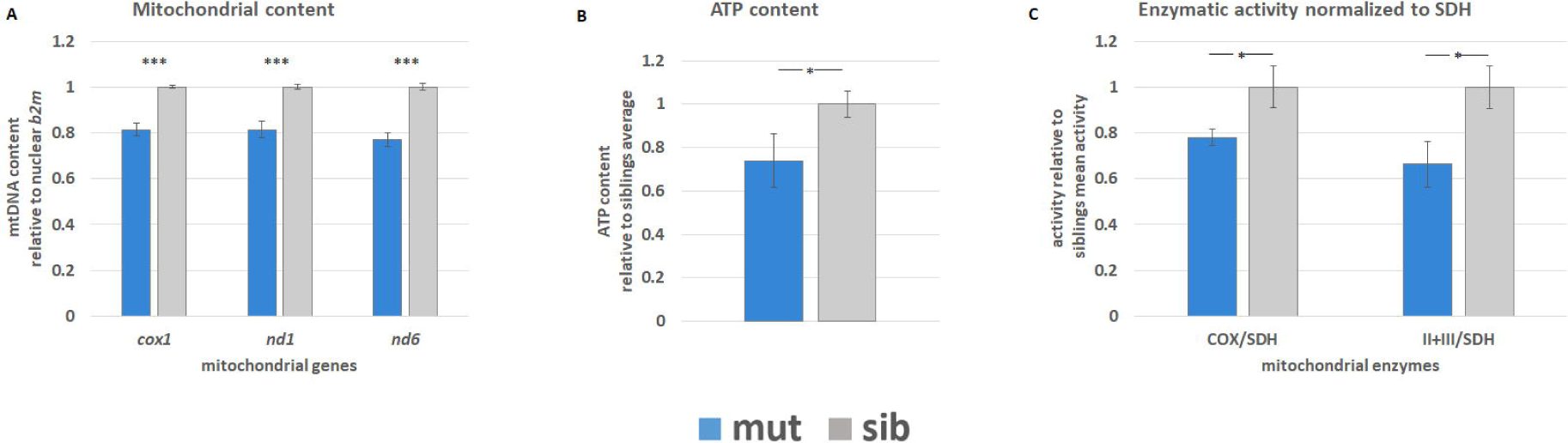
Mitochondrial characterization and ATP content of *atad3*-null larvae. Mutants in blue, siblings in grey. (A) mtDNA content comparison between pools of mutant embryos and their healthy siblings (5dpf); 3 biological replicates of pools (3-8 mutants each, 6-11 siblings each). (B) ATP content in the heads of 6dpf embryos, relative to average sib ATP content measured on the same day; four biological replicates, two from each day. 3 heads in each sample. (C) Enzymatic activity of COX and complex II+III normalized to SDH activity relative to the average enzymatic activity of siblings. Measured from 3 biological replicates, 7-9 heads of 5 dpf larvae in each sample. Error bars indicate standard error of means. Asterisks represent levels of significance (*p<0.05, **p<0.01, ***p<0.001).

### 3.6. ATP content is decreased in heads of mutant larvae compared to siblings

Based on the results of in situ hybridization which demonstrated that *atad3* expression levels was highest in the head, and therefore we determined ATP content in this region. Analysis of mutant 6 dpf larvae relative to the average ATP content in sibling heads collected on the same day revealed a 26% reduction (*p* < 0.05) in ATP content (Fig. 2B).

### 3.7. COX, complex II+III activities are decreased in heads of mutant larvae compared to siblings

In accord with high *atad3* expression levels in the head region, we assesed cytochrome *c* oxidase (COX) and mitochondrial complex II+III activity in heads of mutant larvae. Compared to the average siblings’ activity, COX/SDH activity at 5 dpf was reduced to 78%, and complex II+III/SDH activity was reduced to 66% (*p* < 0.05) (Fig. 2C). Notably, SDH is encoded by nuclear genes only, thus our results indicate a decreased activity of the mtDNA encoded respiratory chain complexes in mutant larvae.

### 3.8. *atad3*-null larvae show decreased locomotor activity

Locomotor assays reveal an overall reduction in movement in the mutant larvae (Fig. 3A). The average distance moved by the siblings was 4.3-fold higher than the mutant larvae (an average of 6.99 m/hour compared to 1.62 m/hour, *p* < 0.001) (Fig. 3B); the difference was greater during light conditions (Fig. 3A). The mutant larvae also had an attenuated immediate reaction to tap (Fig. 3C,D). The average distance moved in the 0.5 second following tap in the mutants group was 69% compared to siblings (an average of 0.497 cm and 0.716 cm, respectively; *p* = 0.012). The mutant larvae also had a reduced light-off immediate reaction (Fig. 3 E,F) of 3.5% compared to siblings (an average of 0.021 cm compared to 0.612 cm moved in the 0.5 second following light-off, respectively; *p* < 0.001) with a comparable movement rate in the following minutes (*p* = 0.67) (Fig.3E,G).

**Figure 3.**
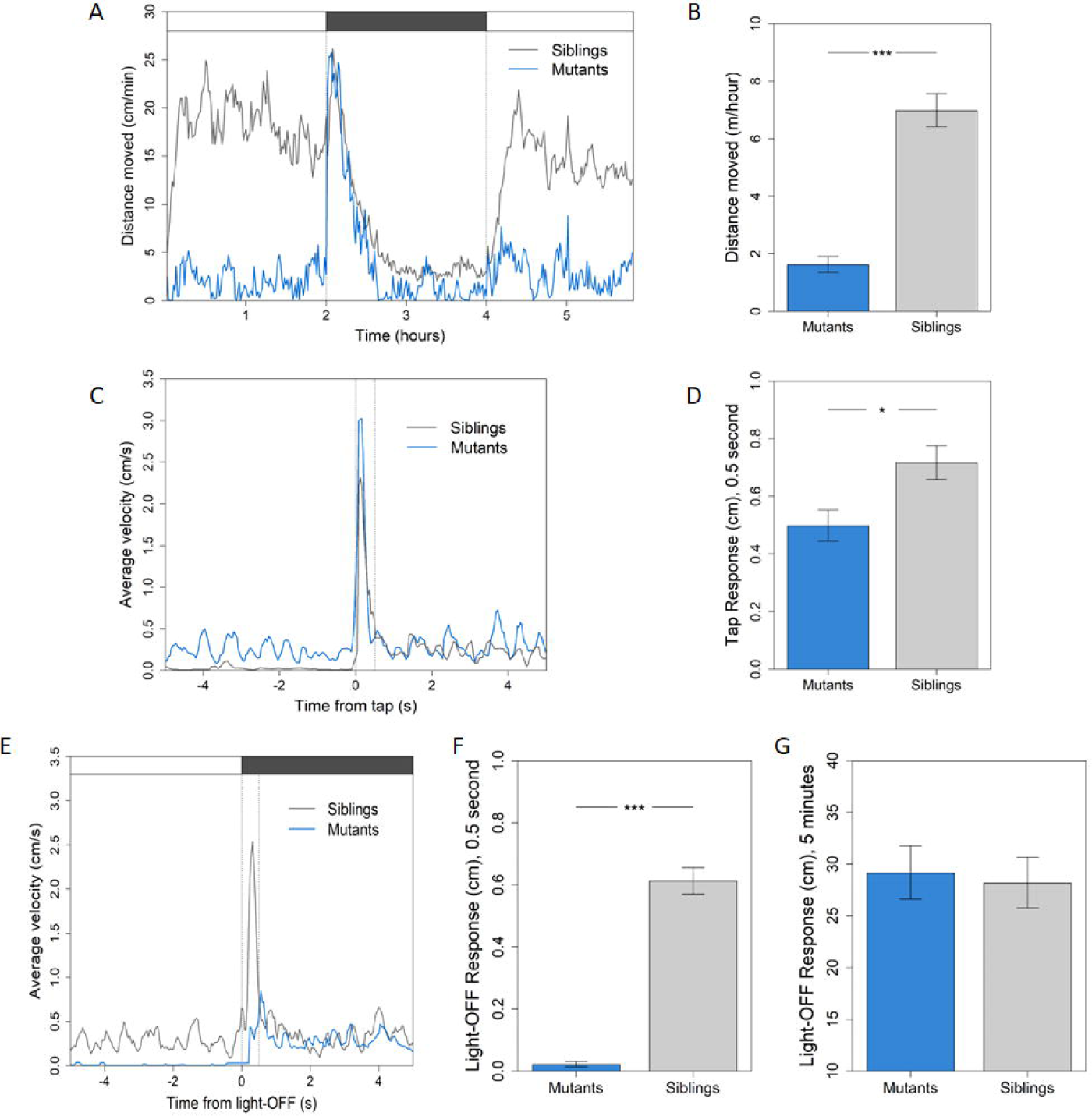
Locomotor activity of *atad3*-null larvae at 5 dpf and 6 dpf is reduced. Mutants in blue, siblings in grey. (A) Activity over time at 5 dpf: average distance moved per minute over 6 hours (light-dark-light cycle, 2 hours each). Upper panel indicates light/dark conditions. (B) Average distance moved over 6 hours is 4.3-times lower in the mutant group compared to siblings at 5 dpf (6.99 m/hour and to 1.62 m/hour, *p* < 0.001). (C-D) Response to tap in mutants compared to siblings, average of 5 and 6 dpf: (C) average velocity of mutant larvae immediately after tap (0 seconds on the X axis) is lower than siblings. (D) Average distance moved €n the first 0.5 sec following tap is 1.4-times lower in mutant larvae compared to siblings (0.497 cm and 0.716 cm, respectively; *p* = 0.012). (E-G) Light-off response in mutant larvae compared to siblings, average of 5 and 6 dpf: € average velocity before and after light-off (at 0 seconds on the X axis). (F) Initial light-off response: average distance in the first o.5 sec following light-off is 29-times lower in mutants compared to siblings (0.021 cm and 0.612 cm, respectively; *p* < 0.001). (G) Average distance moved over 5 minutes after light-off switch shows no difference between mutant and sibling activity (*p* = 0.67). Asterisks represent levels of significance (*p<0.05, **p<0.01, ***p<0.001).

### 3.9. Mutant embryos show significant differential expression of specific biological pathways

To enhance our understanding of the biological pathways perturbed in the mutant embryos, we undertook RNA-seq of homozygous mutant versus their sibling homozygous WT embryos. Siblings were used to minimize batch-specific and timing biases. Extracting DNA and RNA from single embryos enabled individual genotyping followed by pooling of RNA from WT siblings as opposed to all siblings, WT and heterozygotes, providing a significant variance between the two groups (Supplemental Fig. 3). We identified a total of 2109 genes that were differentially expressed in *atad3-* null zebrafish, consisting of 1261 downregulated and 848 upregulated genes (Supplemental Table 2). Gene set enrichment analysis (GSEA) was applied to the whole differential expression data (see Methods section 2.15), and revealed downregulation of tricarboxylic acid cycle, pyruvate metabolism, oxidative phosphorylation, ascorbate and aldarate metabolism, fatty acid metabolism, biosynthesis, oxidation and degradation, and nucleoid dysfunction – all correlating well with mitochondrial dysfunction. Some pathways were shown to be upregulated, including apoptosis, insulin signaling, adipocytokine signaling and Fas pathway and stress induction of heat shock protein regulation. Innate immune genes linked to interferon signaling are also upregulated. Another gene group that was entirely upregulated was the cytosolic aminoacyl-tRNA synthetases (aaRS genes) (Fig.4A). These data support the finding of mitochondrial dysfunction related to known *atad3* pathways in the *atad3*-null zebrafish embryos, as well as to mitochondrial stress and possibly to ER stress. Expression of sample genes from selected groups are shown in heat-map (Fig. 4B), and GSEA plots of four selected gene sets are shown in Fig. 4C. GSEA plots of additional gene sets from Fig. 4A can be found in Supplemental Fig. 4.

**Figure 4.**
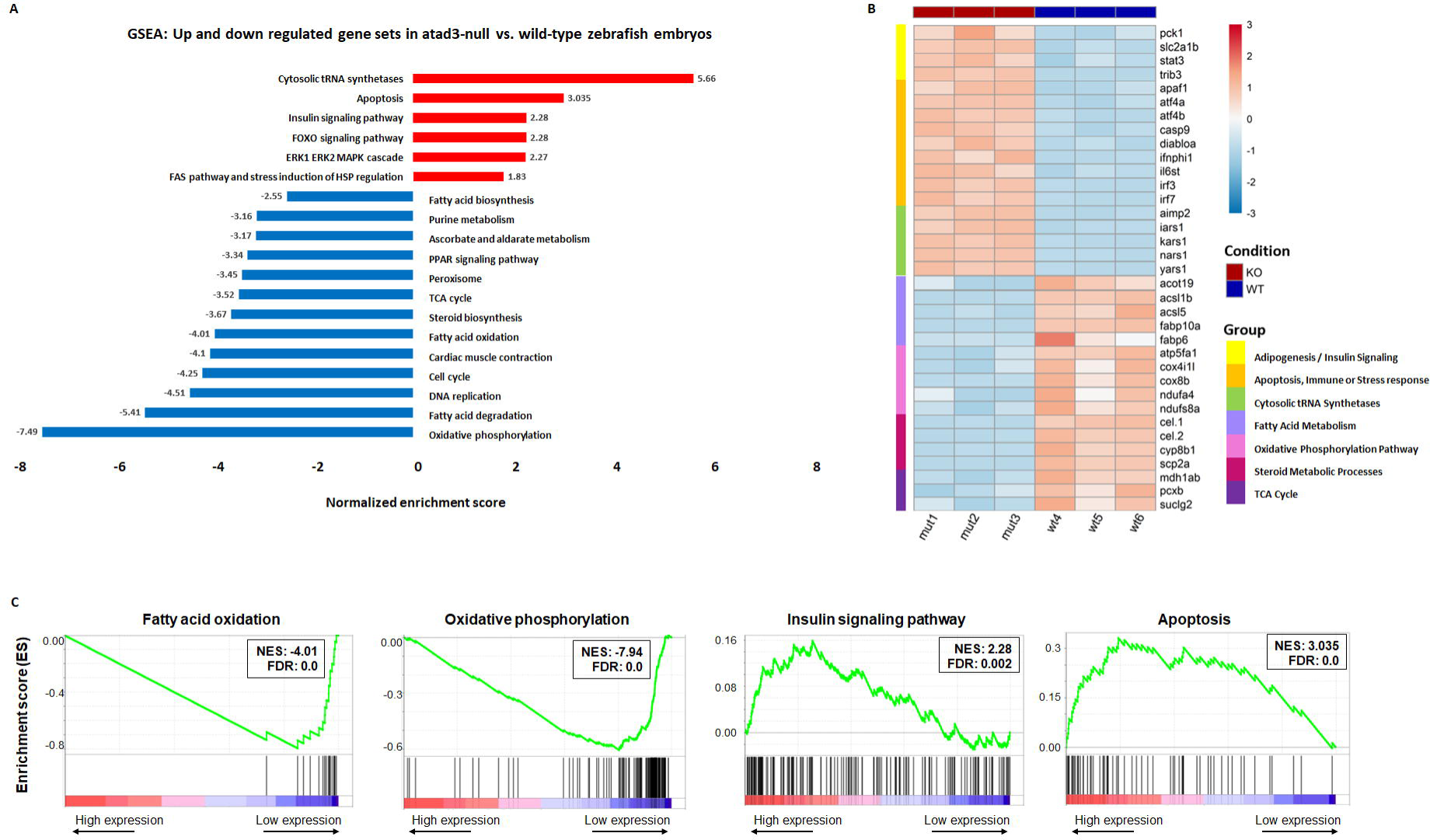
RNA-seq comparing mut 3 dpf embryos to their WT siblings. (A) Selected pathways that were downregulated (blue) or upregulated (red) in the *atad3*-null embryos compared to WT. (B) Heat-map representing the expression of selected genes from selected gene groups in WT vs. mutant embryo pools. (C) Selected GSEA plots showing gene sets that are downregulated or upregulated in KO vs. control. NES: normalized enrichment score; FDR: false discovery rate (corrected *p* value).

To determine whether the differential regulation was specific to 3 dpf, or whether it represented a more global trend of *atad3* mutant vs. WT embryos, we chose several genes for validation by RT-qPCR at both 3 dpf and 5 dpf. According to biological pathways, these included cytosolic tRNA synthetases (*kars1, yars1, lars1*), oxidative phosphorylation (*cox8b, atp5fa1*), and fatty acid metabolism (*acsl5, acot19, fabp10a*) (Fig. 5A). The same trend was confirmed at 5 dpf for all genes tested (Fig. 5B).

**Figure 5.**
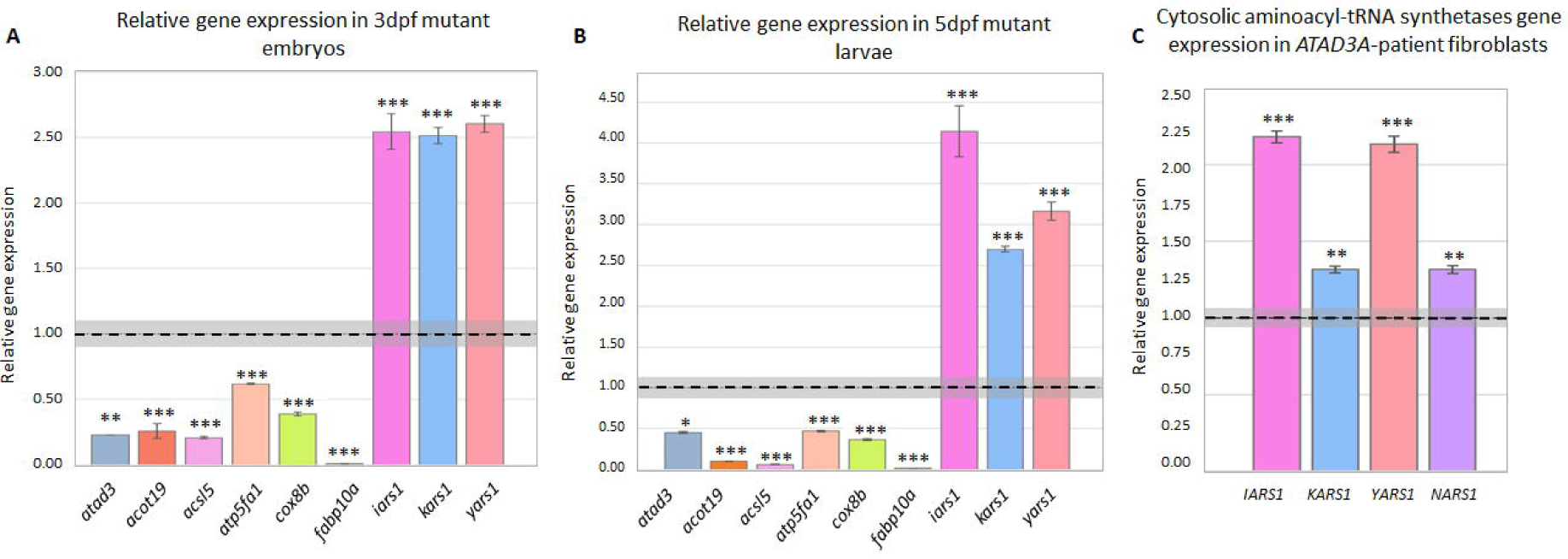
RT-qPCR validations for RNA-seq results. (A,B) RT-qPCR validation of differential expression by major gene groups: (A) at 3dpf. Gene expression in a pool of 20 mutants is normalized to *gapdh* and relative to expression in a pool of 15 siblings, marked by a black dashed line, with siblings average standard error of means of 3 technical replicates in grey. Error bars indicate mutant pool’s standard error of means of 3 technical replicates. (B) at 5dpf. Gene expression in a pool of 9 mutants is normalized to *gapdh* and relative to expression in a pool of 9 WT siblings, marked by a black dashed line, with WT average standard error of means of 3 technical replicates in grey. Error bars indicate mutant pool’s standard error of means of 3 technical replicates. (C) aaRS genes upregulated in fibroblasts derived from an individual with the monoallelic, dominant negative *ATAD3A* variant. Gene expression is normalized to *GAPDH* and relative to expression in control fibroblasts (marked by a black line, with the average standard error of means of 3 technical replicates in grey. Error bars indicate standard error of means in 3 technical replicates of affected fibroblasts. Asterisks represent levels of significance (*p<0.05, **p<0.01, ***p<0.001).

### 3.10. Global rise in cytosolic tRNA synthetases is consistent in fibroblasts derived from an *ATAD3A* patient

To evaluate whether the global rise in cytosolic tRNA synthetases was unique to the zebrafish model or indicative of *ATAD3A* mutant-induced expression changes, RT-qPCR was performed on cDNA derived from fibroblasts of an affected individual with a *de novo* heterozygous *ATAD3A* variant [NM_001170535.3(ATAD3A): c.1582C>T; p.(Arg528Trp)]. This individual was described in detail in Harel et al., 2016 (3). The expression of four randomly chosen cytosolic aminoacyl-tRNA synthetases (*YARS1, IARS1, NARS1, KARS1*) was upregulated in the patient compared to control fibroblasts, consistent with the *atad3*-KO model (Fig. 5C).

## 4. DISCUSSION

Through phenotyping, transcriptional profiling and mitochondrial functional assays, we established a zebrafish *atad3*-null model and its relevance for studying gene function and human *ATAD3A*-related disorders. The observed phenotype, affecting the brain, eyes, heart and muscles, aligns with previous reports in affected individuals (3). Moreover, the reduction in mitochondrial activity as measured by respiratory chain complexes (complex II+III, COX) activity and a reduction in ATP content, reflect a consistent manifestation of mitochondrial disorders (43). While our results indicate a reduction in mtDNA content, further investigations are required to differentiate between defective mtDNA replication and reduced mitochondrial mass (44). Nevertheless, decreased mtDNA content is in accord with decreased mtDNA-dependent activities (II+III and COX) relative to the mtDNA-independent activity of SDH, which is solely encoded in the nuclear genome. With respect to *in vivo* functional manifestations, the diminished locomotor activity of *atad3*-null larvae, both in resting conditions and after stimulation, indicates a motor and/or neuronal impairment. Survival in the mutant embryos is reduced, similarly to the reported human cases (3,19–21,45). The hatching of both mutants and siblings was within the normal time range of 48-72hpf (46). However, hatching of mutants tended to be later within this timeframe. It is unclear whether this observation is indicative of delayed development or muscle weakness (46)(47).

Transcriptional profiling of the *atad3*-null embryos indicated down-regulation of mitochondrial and metabolic pathways known to be connected to human *ATAD3A*, including oxidative phosphorylation, fatty-acids and steroids biosynthesis and metabolism (1,48). Major interferon signaling genes were upregulated, consistent with previous reports in human studies (16). Our study also suggested novel research directions by revealing a specific and global rise in all cytosolic aminoacyl tRNA synthetases (aaRS genes) in *atad3*-null embryos. Other translation pathways and components were not overexpressed, such that this rise could not be attributed to a general non-specific upregulation of translation in the mutant embryos. Apart from the main role of these genes in amino acid biosynthesis, aaRS genes are also known to have non-canonical functions unique to the different genes, related to regulation of various cellular processes including immune response, gene expression regulation, RNA splicing, tumorigenesis and more (49–51). However, the upregulation of all cytosolic aaRS genes suggested a link to their canonical role. Recently, *ATAD3A* was shown to stabilize GRP78 and thus suppress ER stress in colorectal cancer, contributing to chemoresistance (13). The ER stress response may be mediated by three pathways: IRE1 (52), PERK (53), and ATF6 (54). We essentially ruled out involvement of the IRE1 pathway in the context of *atad3* deficiency, since *XBP1* splicing was not altered in the raw transcriptome data in *atad3*-null embryos compared to siblings (results not shown) as expected for this pathway (52). The PERK pathway involves upregulation of *ATF4* (53), which has interestingly been shown to upregulate aaRS genes (55), and was significantly upregulated in the mutant zebrafish. Taken together, this suggests that the ER stress response in the context of *atad3* deficiency may be mediated by this pathway.

A similar upregulation in aaRS genes mediated by ATF4 was previously reported, triggered by mitochondrial stress induced by various effectors on mammalian cells (56) ATF4 is also active in a non-canonical mitochondrial stress response (mitochondrial unfolded protein response, UPR^mt^) [56]. Nonetheless, we did not observe upregulation of other genes associated with the UPR^mt^ response in the mutant zebrafish. Therefore, we hypothesize that the upregulation of *ATF4* and aaRS genes is mediated by mitochondrial stress and the integrated stress response (IRS), with possible influence from ER stress.

Our transcriptome results also suggest the *atad3-*null zebrafish embryos have a higher level of apoptosis. Furthermore, it shows elevated insulin signaling, adipogenesis and adipocytokine signaling pathways, all pointing towards insulin resistance and related to the known *ATAD3A* effect on fatty-acid metabolism. Mitochondrial dysfunction and specifically ER-mitochondria contact site dysfunction have been shown to lead to insulin resistance (57–59). Interestingly, a ketogenic diet was previously shown to improve neurological symptoms in patients with *ATAD3A* variants. While one of the variants mentioned (p.Thr84Met, occurring as a biallelic variant in one patient (60) and in trans to a deletion in another patient (61)) was not proven pathogenic, the phenotype of the patients fits the diagnosis of *ATAD3A*-disorder, which was improved by a ketogenic diet (60,61).

The human paralogs *ATAD3A* and *ATAD3B* are highly similar, yet have some distinct functions. Importantly, *ATAD3A* is the only ubiquitously expressed paralog in human, although *ATAD3B* can have a role in disease (10,62,63). While the lack of *atad3* gene paralogs in zebrafish introduces a potential difference from the human context, the model’s resemblance to human *ATAD3A-*associated diseases suggests its utility in advancing our understanding of these disorders.

In conclusion, transcriptome analysis of mutant versus WT embryos underscored the importance of mitochondrial dysfunction in the pathogenesis of *ATAD3A-*related disorders and demonstrated an unexpected elevation of cytosolic aaRS genes, secondary to mitochondrial stress and possibly to elevated ER stress. This zebrafish model opens avenues for investigating therapeutic interventions and further exploring the underlying mechanisms of *ATAD3A-*related disorders.

## DECLARATIONS

### Ethics approval and consent to participate

Patient fibroblasts were obtained under research protocol H-29697, approved by the Institutional Review Board (IRB) at Baylor College of Medicine (BCM), and previously published (3).

### Consent for publication

Informed consent was obtained for research and publication.

### Availability of data and material

RNAseq data was submitted to Gene Expression Omnibus (GEO): https://www.ncbi.nlm.nih.gov/geo/query/acc.cgi?acc=GSE236968

## Authors’ contributions

SE and TH designed the experiments and wrote the manuscript. SE, NR, SRB, OH conducted the experiments. AS oversaw mitochondrial assays. YW and ZB conducted and analyzed locomotor assays. IP and HB provided bioinformatic support and analyzed data. SYD, AS and AI designed experiments and contributed to manuscript writing. All authors reviewed and approved the manuscript. **Acknowledgements.** Not applicable.

## List of abbreviations

aaRS: aminoacyl-tRNA synthetases
ADP: adenosine triphosphate
COX: cytochrome *c* oxidase dpf – days post-fertilization
ER: endoplasmic reticulum
GOBP: Gene Ontology Biological Process
GSEA: Gene Set Enrichment Analysis
KEGG: Kyoto Encyclopedia of Genes and Genomes
KO: knockout
MAM: mitochondrial associated membranes
mtDNA: mitochondrial DNA
NAFLD: nonalcoholic fatty liver disease
NAHR: nonallelic homologous recombination
SDH: succinate dehydrogenase
WK: WikiPathways
WT: wild type

## REFERENCES

1. Arguello T, Peralta S, Antonicka H, Gaidosh G, Diaz F, Tu YT, et al. ATAD3A has a scaffolding role regulating mitochondria inner membrane structure and protein assembly. Cell Rep. 2021;37(12).

2. Gilquin B, Taillebourg E, Cherradi N, Hubstenberger A, Gay O, Merle N, et al. The AAA + ATPase ATAD3A Controls Mitochondrial Dynamics at the Interface of the Inner and Outer Membranes. Mol Cell Biol. 2010;30(8).

3. Harel T, Yoon WH, Garone C, Gu S, Coban-Akdemir Z, Eldomery MK, et al. Recurrent De Novo and Biallelic Variation of ATAD3A, Encoding a Mitochondrial Membrane Protein, Results in Distinct Neurological Syndromes. Am J Hum Genet. 2016;99(4).

4. He J, Cooper HM, Reyes A, Di Re M, Sembongi H, Gao J, et al. Mitochondrial nucleoid interacting proteins support mitochondrial protein synthesis. Nucleic Acids Res. 2012;40(13).

5. Ishihara T, Ban-Ishihara R, Ota A, Ishihara N. Mitochondrial nucleoid trafficking regulated by the inner-membrane AAA-ATPase ATAD3A modulates respiratory complex formation. Proc Natl Acad Sci U S A. 2022;119(47).

6. Rone MB, Midzak AS, Issop L, Rammouz G, Jagannathan S, Fan J, et al. Identification of a dynamic mitochondrial protein complex driving cholesterol import, trafficking, and metabolism to steroid hormones. Molecular Endocrinology. 2012;26(11).

7. Chen L, Li Y, Sottas C, Lazaris A, Petrillo SK, Metrakos P, et al. Loss of mitochondrial ATPase ATAD3A contributes to nonalcoholic fatty liver disease through accumulation of lipids and damaged mitochondria. J Biol Chem. 2022;298(6).

8. Zhao Y, Hu D, Wang R, Sun X, Ropelewski P, Hubler Z, et al. ATAD3A oligomerization promotes neuropathology and cognitive deficits in Alzheimer’s disease models. Nat Commun. 2022;13(1).

9. Gerhold JM, Cansiz-Arda S, Lohmus M, Engberg O, Reyes A, Van Rennes H, et al. Human Mitochondrial DNA-Protein Complexes Attach to a Cholesterol-Rich Membrane Structure. Sci Rep. 2015;5.

10. Baudier J. ATAD3 proteins: brokers of a mitochondria–endoplasmic reticulum connection in mammalian cells. Biological Reviews. 2018;93(2).

11. Watanabe S, Horiuchi M, Murata Y, Komine O, Kawade N, Sobue A, et al. Sigma-1 receptor maintains ATAD3A as a monomer to inhibit mitochondrial fragmentation at the mitochondria-associated membrane in amyotrophic lateral sclerosis. Neurobiol Dis. 2023;179.

12. Lang L, Loveless R, Teng Y. Emerging links between control of mitochondrial protein atad3a and cancer. Vol. 21, International Journal of Molecular Sciences. 2020.

13. Huang KCY, Chiang SF, Yang PC, Ke TW, Chen TW, Lin CY, et al. ATAD3A stabilizes GRP78 to suppress ER stress for acquired chemoresistance in colorectal cancer. J Cell Physiol. 2021;236(9).

14. Xie XQ, Yang Y, Wang Q, Liu HF, Fang XY, Li CL, et al. Targeting ATAD3A-PINK1-mitophagy axis overcomes chemoimmunotherapy resistance by redirecting PD-L1 to mitochondria. Cell Res. 2023;33(3).

15. Di Rienzo M, Romagnoli A, Ciccosanti F, Refolo G, Consalvi V, Arena G, et al. AMBRA1 regulates mitophagy by interacting with ATAD3A and promoting PINK1 stability. Autophagy. 2022;18(8).

16. Lepelley A, Mina E Della, van Nieuwenhove E, Waumans L, Fraitag S, Rice GI, et al. Enhanced cGAS-STING–dependent interferon signaling associated with mutations in ATAD3A. Journal of Experimental Medicine. 2021;218(10).

17. Merle N, Féraud O, Gilquin B, Hubstenberger A, Kieffer-Jacquinot S, Assard N, et al. ATAD3B is a human embryonic stem cell specific mitochondrial protein, re-expressed in cancer cells, that functions as dominant negative for the ubiquitous ATAD3A. Mitochondrion. 2012;12(4).

18. Desai R, Frazier AE, Durigon R, Patel H, Jones AW, Rosa ID, et al. ATAD3 gene cluster deletions cause cerebellar dysfunction associated with altered mitochondrial DNA and cholesterol metabolism. Brain. 2017;140(6).

19. Yap ZY, Park YH, Wortmann SB, Gunning AC, Ezer S, Lee S, et al. Functional interpretation of ATAD3A variants in neuro-mitochondrial phenotypes. Genome Med. 2021;13(1).

20. Gunning AC, Strucinska K, Muñoz Oreja M, Parrish A, Caswell R, Stals KL, et al. Recurrent De Novo NAHR Reciprocal Duplications in the ATAD3 Gene Cluster Cause a Neurogenetic Trait with Perturbed Cholesterol and Mitochondrial Metabolism. Am J Hum Genet. 2020;106(2).

21. Frazier AE, Compton AG, Kishita Y, Hock DH, Welch AE, Amarasekera SSC, et al. Fatal perinatal mitochondrial cardiac failure caused by recurrent de novo duplications in the ATAD3 locus. Med (N Y). 2021 Jan 15;2(1):49–73.

22. Cooper HM, Yang Y, Ylikallio E, Khairullin R, Woldegebriel R, Lin KL, et al. ATpase-deficient mitochondrial inner membrane protein ATAD3a disturbs mitochondrial dynamics in dominant hereditary spastic paraplegia. Hum Mol Genet. 2017;26(8).

23. Fichi G, Naef V, Barca A, Longo G, Fronte B, Verri T, et al. Fishing in the cell powerhouse: Zebrafish as a tool for exploration of mitochondrial defects affecting the nervous system. Vol. 20, International Journal of Molecular Sciences. 2019.

24. Labun K, Montague TG, Krause M, Torres Cleuren YN, Tjeldnes H, Valen E. CHOPCHOP v3: Expanding the CRISPR web toolbox beyond genome editing. Nucleic Acids Res. 2019;47(W1).

25. Meeker ND, Hutchinson SA, Ho L, Trede NS. Method for isolation of PCR-ready genomic DNA from zebrafish tissues. Biotechniques. 2007;43(5).

26. Zhu X, Xu Y, Yu S, Lu L, Ding M, Cheng J, et al. An efficient genotyping method for genome-modified animals and human cells generated with CRISPR/Cas9 system. Sci Rep. 2014;4.

27. Saada A, Bar-Meir M, Belaiche C, Miller C, Elpeleg O. Evaluation of enzymatic assays and compounds affecting ATP production in mitochondrial respiratory chain complex I deficiency. Anal Biochem. 2004;335(1).

28. Ben-Meir A, Yahalomi S, Moshe B, Shufaro Y, Reubinoff B, Saada A. Coenzyme Q-dependent mitochondrial respiratory chain activity in granulosa cells is reduced with aging. Fertil Steril. 2015;104(3).

29. Noldus LPJJ, Spink AJ, Tegelenbosch RAJ. EthoVision: A versatile video tracking system for automation of behavioral experiments. Behavior Research Methods, Instruments, and Computers. 2001;33(3).

30. Burgess HA, Granato M. Modulation of locomotor activity in larval zebrafish during light adaptation. Journal of Experimental Biology. 2007;210(14).

31. Sakov A, Golani I, Lipkind D, Benjamini Y. High-throughput data analysis in behavior genetics. Annals of Applied Statistics. 2010;4(2).

32. Emran F, Rihel J, Adolph AR, Wong KY, Kraves S, Dowling JE. OFF ganglion cells cannot drive the optokinetic reflex in zebrafish. Proc Natl Acad Sci U S A. 2007;104(48).

33. Benjamini Y, Hochberg Y. Controlling the False Discovery Rate: A Practical and Powerful Approach to Multiple Testing. Journal of the Royal Statistical Society: Series B (Methodological). 1995;57(1).

34. Kuznetsova A, Brockhoff PB, Christensen RHB. lmerTest Package: Tests in Linear Mixed Effects Models. J Stat Softw. 2017;82(13).

35. Lenth R. emmeans: estimated marginal means, aka least-squares means. R Package version 145. 2020;

36. Martin M. Cutadapt removes adapter sequences from high-throughput sequencing reads. EMBnet J. 2011;17(1).

37. Kim D, Pertea G, Trapnell C, Pimentel H, Kelley R, Salzberg SL. TopHat2: Accurate alignment of transcriptomes in the presence of insertions, deletions and gene fusions. Genome Biol. 2013;14(4).

38. Anders S, Pyl PT, Huber W. HTSeq-A Python framework to work with high-throughput sequencing data. Bioinformatics. 2015;31(2).

39. Love MI, Huber W, Anders S. Moderated estimation of fold change and dispersion for RNA-seq data with DESeq2. Genome Biol. 2014;15(12).

40. Subramanian A, Tamayo P, Mootha VK, Mukherjee S, Ebert BL, Gillette MA, et al. Gene set enrichment analysis: A knowledge-based approach for interpreting genome-wide expression profiles. Proc Natl Acad Sci U S A. 2005;102(43).

41. Kuleshov M V., Jones MR, Rouillard AD, Fernandez NF, Duan Q, Wang Z, et al. Enrichr: a comprehensive gene set enrichment analysis web server 2016 update. Nucleic Acids Res. 2016;44(1).

42. Waters ER, Bezanilla M, Vierling E. ATAD3 Proteins: Unique Mitochondrial Proteins Essential for Life in Diverse Eukaryotic Lineages. Plant Cell Physiol. 2023 Oct 19;

43. Gajewski CD, Yang L, Schon EA, Manfredi G. New insights into the bioenergetics of mitochondrial disorders using intracellular ATP reporters. Mol Biol Cell. 2003;14(9).

44. Malik AN, Czajka A. Is mitochondrial DNA content a potential biomarker of mitochondrial dysfunction? Mitochondrion. 2013;13(5).

45. Peralta S, González-Quintana A, Ybarra M, Delmiro A, Pérez-Pérez R, Docampo J, et al. Novel ATAD3A recessive mutation associated to fatal cerebellar hypoplasia with multiorgan involvement and mitochondrial structural abnormalities. Mol Genet Metab. 2019;128(4).

46. Kimmel CB, Ballard WW, Kimmel SR, Ullmann B, Schilling TF. Stages of embryonic development of the zebrafish. Developmental Dynamics. 1995;203(3).

47. Ahmad F, Ali S, Richardson MK. Effect of pesticides and metals on zebrafish embryo development and larval locomotor activity. bioRxiv. 2020;

48. Issop L, Fan J, Lee S, Rone MB, Basu K, Mui J, et al. Mitochondria-Associated membrane formation in hormone-stimulated leydig cell steroidogenesis: Role of ATAD3. Endocrinology. 2015;156(1).

49. Turvey AK, Horvath GA, Cavalcanti ARO. Aminoacyl-tRNA synthetases in human health and disease. Vol. 13, Frontiers in Physiology. 2022.

50. Wakasugi K, Yokosawa T. Non-canonical functions of human cytoplasmic tyrosyl-, tryptophanyl-and other aminoacyl-tRNA synthetases. In: Enzymes. 2020.

51. Ivanov KA, Moor NA, Lavrik OI. Non-canonical functions of aminoacyl-tRNA synthetases. Biochemistry (Mosc). 2000 Aug;65(8):888–97.

52. Cox JS, Shamu CE, Walter P. Transcriptional induction of genes encoding endoplasmic reticulum resident proteins requires a transmembrane protein kinase. Cell. 1993;73(6).

53. Harding HP, Zhang Y, Ron D. Protein translation and folding are coupled by an endoplasmic-reticulum-resident kinase. Nature. 1999;397(6716).

54. Haze K, Yoshida H, Yanagi H, Yura T, Mori K. Mammalian transcription factor ATF6 is synthesized as a transmembrane protein and activated by proteolysis in response to endoplasmic reticulum stress. Mol Biol Cell. 1999;10(11).

55. Gonen N, Meller A, Sabath N, Shalgi R. Amino Acid Biosynthesis Regulation during Endoplasmic Reticulum Stress Is Coupled to Protein Expression Demands. iScience. 2019;19.

56. Quirós PM, Prado MA, Zamboni N, D’Amico D, Williams RW, Finley D, et al. Multi-omics analysis identifies ATF4 as a key regulator of the mitochondrial stress response in mammals. Journal of Cell Biology. 2017;216(7).

57. Rieusset J. Contribution of mitochondria and endoplasmic reticulum dysfunction in insulin resistance: Distinct or interrelated roles? Vol. 41, Diabetes and Metabolism. 2015.

58. Arruda AP, Pers BM, Parlakgül G, Güney E, Inouye K, Hotamisligil GS. Chronic enrichment of hepatic endoplasmic reticulum-mitochondria contact leads to mitochondrial dysfunction in obesity. Nat Med. 2014;20(12).

59. Koliaki C, Roden M. Hepatic energy metabolism in human diabetes mellitus, obesity and non-alcoholic fatty liver disease. Vol. 379, Molecular and Cellular Endocrinology. 2013.

60. Madhoun A Al, Alnaser F, Melhem M, Nizam R, Al-Dabbous T, Al-Mulla F. Ketogenic diet attenuates cerebellar atrophy progression in a subject with a biallelic variant at the ATAD3A locus. Application of Clinical Genetics. 2019;12.

61. Chen Y, Rong S, Luo H, Huang B, Hu F, Chen M, et al. Ketogenic diet attenuates refractory epilepsy of Harel-Yoon syndrome with ATAD3A variants: a case report and review of literature. Pediatr Neurol. 2023;

62. Shu L, Hu C, Xu M, Yu J, He H, Lin J, et al. ATAD3B is a mitophagy receptor mediating clearance of oxidative stress-induced damaged mitochondrial DNA. EMBO J. 2021;40(8).

63. Yanovsky-Dagan S, Frumkin A, Lupski JR, Harel T. CRISPR/Cas9-induced gene conversion between ATAD3 paralogs. Human Genetics and Genomics Advances. 2022;3(2).

