## Supplemental data for "Transcriptome analysis of *atad3-*null zebrafish embryos elucidates possible disease mechanisms"

4 Department of Neurobiology, The George S. Wise Faculty of Life Sciences, Tel-Aviv University, Tel Aviv, Israel

**SUPPLEMENTAL DATA:**

- **Figure S1. *atad3* expression in zebrafish embryos.**
- **Figure S2. Guide 1 (gd1)-targeted CRISPR/Cas9 causes the same effect as gd3-targeted CRISPR/Cas9**
- **Figure S3. Principal component analysis (PCA)**
- **Figure S4. RNA-seq results: differentially expressed pathways**
- **Figure S5. Clustal Omega alignment of *ATAD3* human paralogs with the zebrafish ortholog *atad3***
- **Table S1. Primers and crRNA sequences**
- **Table S2. Differentially expressed genes (attached as a separate excel file)**

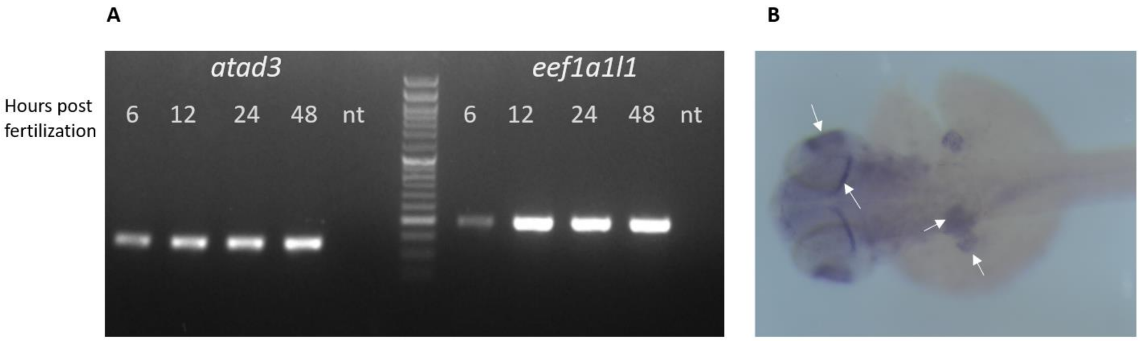


**Figure S1. *atad3* expression in zebrafish embryos.** (A) *atad3* expression in embryos by hours post fertilization (hpf) and nt control (no template) compared to *eef1a1l1* expression (housekeeping gene), RT-PCR; (B) In situ hybridization of *atad3* in a 48hpf zebrafish embryo showing expression (in purple) mainly in the brain, eyes, fin-buds and liver (indicated by white arrows).

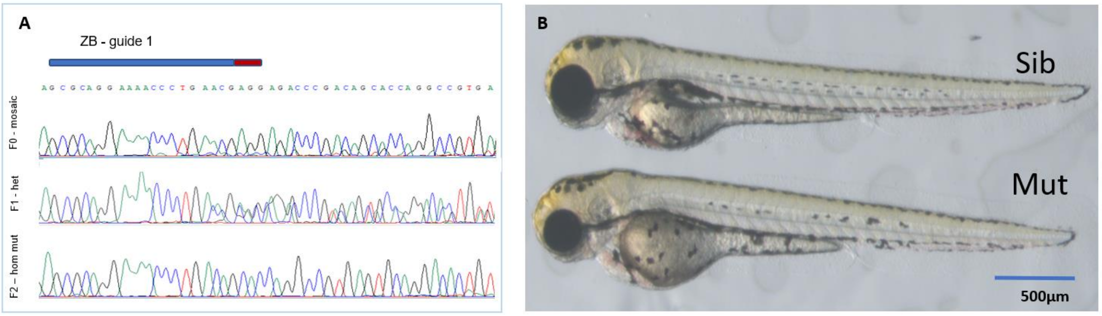


**Figure S2. Guide 1 (gd1)-targeted CRISPR/cas9 causes the same effect as gd3-targeted CRISPR/cas9.** (A) DNA extracted from mutant embryos was sequenced to confirm a homozygous out-of-frame deletion. Upper panel – F0 generation, mosaic (CRISPR/Cas9-injected fish); middle panel – F1 generation (heterozygous); lower panel – homozygous mutant. CRISPR RNA (crRNA) marked by a blue line, PAM sequence marked by a red line. (B) Sib (either WT or heterozygous) vs. mutant embryo at 3dpf. The mutant embryos show the same phenotype as the gd3 mutants.

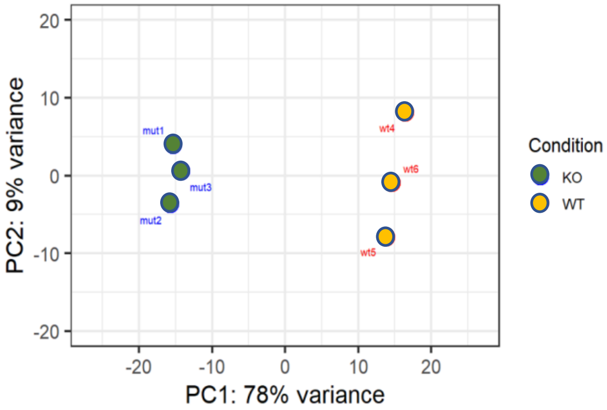


**Figure S3. Principal component analysis (PCA)** indicating global significant difference in RNA expression between WT (in yellow) and mut (in green) embryo pools.

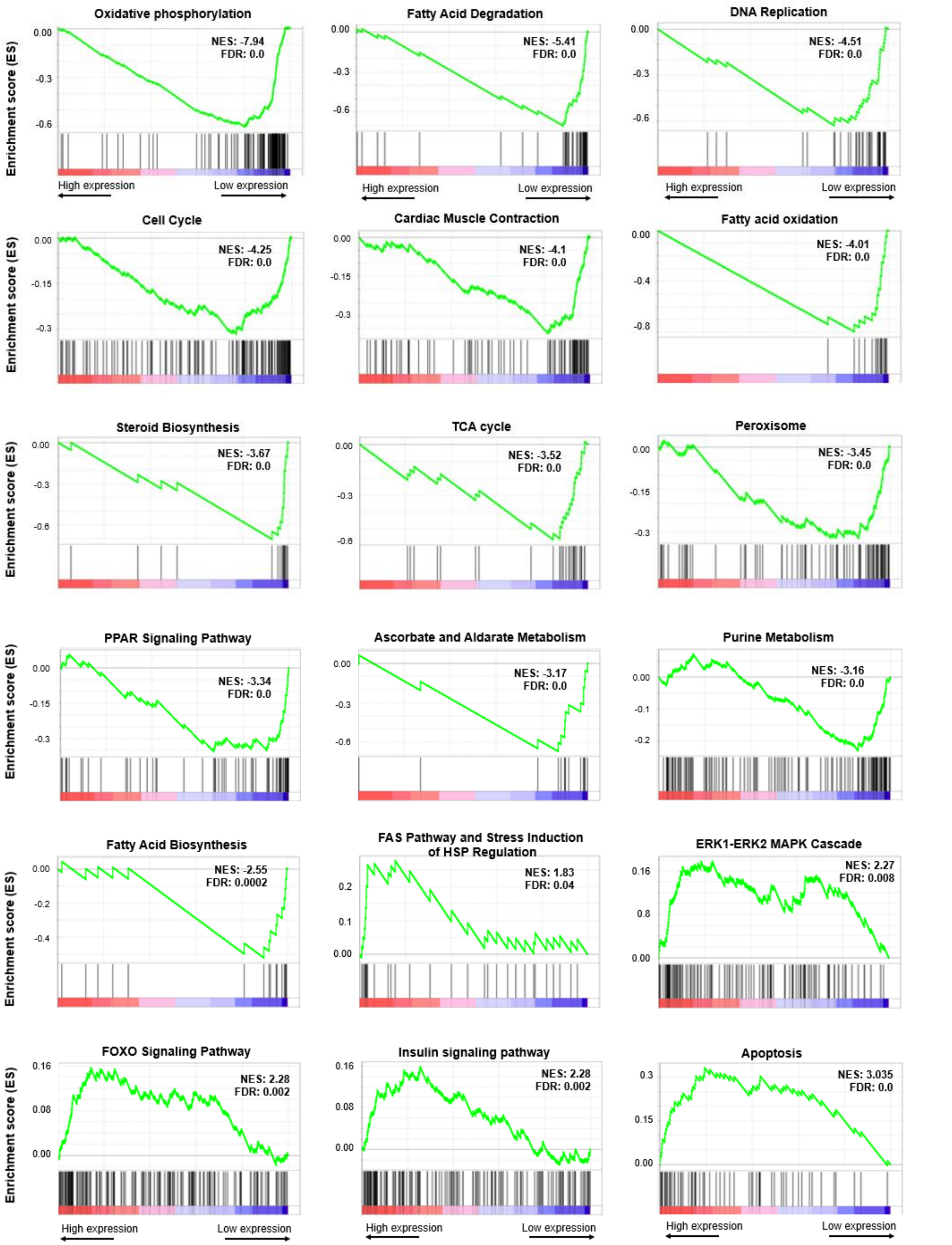


**Figure S4. RNA-seq results: differentially expressed pathways.** Selected GSEA plots showing gene sets that are downregulated or upregulated in KO vs. control. NES: normalized enrichment score; FDR: false discovery rate (corrected *p* value).

**Figure S5.** Clustal Omega alignment of *ATAD3* human paralogs with the zebrafish ortholog *atad3*

```
ATAD3A         MSWLFGINKGPKGEGA-GPPPPLPPAQPGAEGGGDRGLGDRPAPKDKWSNFDPTGLERAA    59
ATAD3B         MSWLFGVNKGPKGEGA-GPPPPLPPAQPGAEGGGDRGLGDRPAPKDKWSNFDPTGLERAA    59
atad3          MSWLFGLNKGQSGGPPELPPPPAPPAPPGGSG-------AADKPKDKWSNFDPTGLERAA    53
               ******:*** .*     **** *** **..*           *****************

ATAD3A         KAARELEHSRYAKDALNLAQMQEQTLQLEQQSKLKMRLEALSLLHTLVWAWSLCRAGAVQ    119
ATAD3B         KAARELEHSRYAKEALNLAQMQEQTLQLEQQSKL--------------------------    93
atad3          QAARELDQSRHAKEALDLARMQEQTVQMEHQGKI--------------------------    87
               :*****::**:**:**:**:*****:*:*:*.*:

ATAD3A         TQERLSGSASPEQVPAGECCALQEYEAAVEQLKSEQIRAQAEERRKTLSEETRQHQARAQ    179
ATAD3B         ----------------------KEYEAAVEQLKSEQIRAQAEERRKTLSEETRQHQARAQ    131
atad3          ----------------------KEYEAAVEQLKGEQIRIQADERRKTLNEETRQHQARAQ    125
                                     :**********.**** **:******.***********

ATAD3A         YQDKLARQRYEDQLKQQQLLNEENLRKQEESVQKQEAMRRATVEREMELRHKNEMLRVEA    239
ATAD3B         YQDKLARQRYEDQLKQQQLLNEENLRKQEESVQKQEAMRRATVEREMELRHKNEMLRVET    191
atad3          YQDKLARQRYDDQLRQQTLLNEENLRKQEESVQKQEAMRRATIEHEMDLRHKNEMLRVEA    185
               **********:***:** ************************:*:**:***********:

ATAD3A         EARARAKAERENADIIREQIRLKAAEHRQTVLESIRTAGTLFGEGFRAFVTDWDKVTATV    299
ATAD3B         EARARAKAERENADIIREQIRLKASEHRQTVLESIRTAGTLFGEGFRAFVTDRDKVTATV    251
atad3          ESKARARVERENADIIREQIRLKAAEHRQTVLESIRTAGAVFGEGFRAFISDWDKVTATV    245
               *::***:.****************:**************::********::* *******

ATAD3A         AGLTLLAVGVYSAKNATLVAGRFIEARLGKPSLVRETSRITVLEALRHPIQVSRRLLSRP    359
ATAD3B         AGLTLLAVGVYSAKNATAVTGRFIEARLGKPSLVRETSRITVLEALRHPIQVSRRLLSRP    311
atad3          AGLTLLAAGVYSARNATAVAGRYIEARLGKPSLVRETSRFTVAEALKHPIKVVKRLQSKP    305
               *******.*****:*** *:**:****************:** ***:***:* :** *:*

ATAD3A         QDALEGVVLSPSLEARVRDIAIATRNTKKNRSLYRNILMYGPPGTGKTLFAKKLALHSGM    419
ATAD3B         QDVLEGVVLSPSLEARVRDIAIATRNTKKNRGLYRHILLYGPPGTGKTLFAKKLALHSGM    371
atad3          QDALEGVVLSPPLEERVRDIAIATRNTRQNRGLYRNILMYGPPGTGKTLFAKKLAVHSGM    365
               **.******** ** ************::**.***:**:****************:****

ATAD3A         DYAIMTGGDVAPMGREGVTAMHKLFDWANTSRRGLLLFVDEADAFLRKRATEKISEDLRA    479
ATAD3B         DYAIMTGGDVAPMGREGVTAMHKLFDWANTSRRGLLLFMDEADAFLRKRATEEISKDLRA    431
atad3          DYAIMTGGDVAPMGRDGVTAMHKVFDWAATSRRGLLLFVDEADAFLRKRSTEKISEDLRA    425
               ***************:*******:**** *********:**********:**:**:****

ATAD3A         TLNAFLYRTGQHSNKFMLVLASNQPEQFDWAINDRINEMVHFDLPGQEERERLVRMYFDK    539
ATAD3B         TLNAFLYHMGQHSNKFMLVLASNLPEQFDCAINSRIDVMVHFDLPQQEERERLVRLHFDN    491
atad3          TLNAFLYRTGEQSNKFMLVLASNQPEQFDWAINDRIDEIVNFMLPGPEERERLVRLYFDR    485
               *******: *::*********** ***** ***.**: :*:* **  ********::**.

ATAD3A         YVLKPATEGKQRLKLAQFDYGRKCSEVARLTEGMSGREIAQLAVSWQATAYASEDGVLTE    599
ATAD3B         CVLKPATEGKRRLKLAQFDYGRKCSEVARLTEGMSGREIAQLAVSWQATAYASKDGVLTE    551
atad3          YVLEPATGGRQRLKLAQFDYGQKCSEIAKRVEGMSGREISKLGVAWQAAAYSSEDGVLSE    545
                **:*** *::**********:****:*:* .********::*.*:***:**:*:****:*

ATAD3A         AMMDTRVQDAVQQHQQKMCWLKAEGPGRGDEPSPS-------------------------    634
ATAD3B         AMMDACVQDAVQQYRQKMRWLKAEGPGRGVEHPLSGVQGETLTSWSLATDPSYPCLAGPC    611
atad3          AMIDARVDAAVRQHRQKMDWLHGEGVLDNEGRPVAAETQTKAANMG--FTPPL---KAQE    600
               **:*: *: **:*::*** **:.**   .     :

ATAD3A         -------------------------------------  634
ATAD3B         TFRICSWMGTGLCPGPLSPRMSCGGGRPFCPPGHPLL  648
atad3          EL----------RPLQEVPE-----DSQNKHDGTPV-  621
```

Color codes for *ATAD3A* domains (taken from Unirpot and Waters et al., 2023 [61])

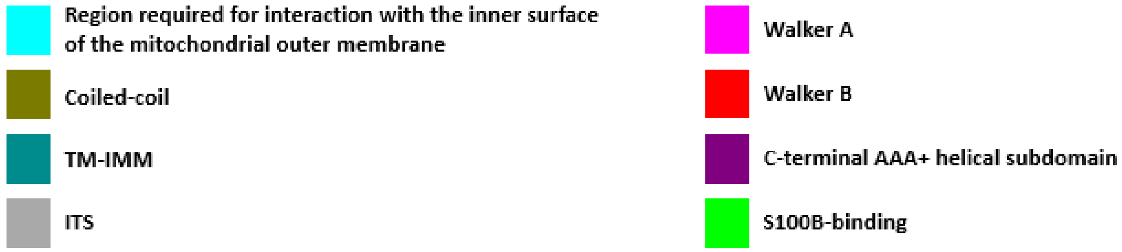


### Table S1. primers and crRNA sequences

| **Primers for semi-quantitative RT-PCR** | | | | |
|---|---|---|---|---|
| **Gene** | **F primer** | **R primer** | **Product size** | **Temp** |
| *atad3* | GGACAAATGGAGCAACTTCG | ATCTGAACCGTCTGCTCCTG | 129 | 60 |
| *eef1a1l1* (housekeeping) | ATCTGATCTACAAATGCGGT | GCATCAATAATGGTGACGTAG | 199 | 59 |
| **Primers for sequencing and genotyping** | | | | |
| **Gene** | **F primer** | **R primer** | **Product size** | **Temp** |
| *atad3* (gd3, for acrylamide genotyping) | GTGTTTTACCTTGATCTTGCCC | AGCATGTGTGACCCCTGTG | 112 | 60 |
| *atad3* (gd1, for acrylamide genotyping) | CAGCTGAAGGGTGAGCAGAT | AACAGTGTCCTGCACTCACG | 100 | 60 |
| *atad3* (gd3, for sequencing) | GTGTTTTACCTTGATCTTGCCC | AGGCGAACTCACACTCTGCT | 168 | 60 |
| *atad3* (gd1, for sequencing) | AACAGTGTCCTGCACTCACG | CAGGAGTACGAGGCAGCAGT | 124 | 60 |
| **Primers for mitochondrial content (qPCR)** | | | | |
| **Gene** | **F primer** | **R primer** | **Product size** | **Temp** |
| *b2m* | CGCCTGAAAACTACGTTCTACAC | ACTTTCGGAGTGGCTGAAAA | 140 | 60 |
| *mt-nd1* | AGCCTACGCCGTACCAGTATT | GTTTCACGCCATCAGCTACTG | 143 | 60 |
| *mt-cox1* | TGGAAACTGACTTGTGCCACT | TCATCCTGTTCCAGCTCCTG | 145 | 60 |
| *mt-nd6* | AGTAGTTGTGGCAGGGGTTG | CTTCAGGAAAAGGCTCGGCT | 146 | 60 |

| RT-qPCR primers | | | | |
|---|---|---|---|---|
| Gene | F primer | R primer | Product size | temp |
| *gapdh* (housekeeping) | CGCTGGCATCTCCCTCAA | TCAGCAACACGATGGCTGTAG | 85 | 60 |
| *atad3* | GGACAAATGGAGCAACTTCG | ATCTGAACCGTCTGCTCCTG | 129 | 60 |
| *kars1* | GTTTCCACCTCCTGACACCT | AAGAAGTCGAGCAGTTGTCCT | 112 | 59 |
| *iars1* | CCATCTGGCATATTCACTCCGG | CCGTTCTCCTTCAGCCACTT | 95 | 60 |
| *yars1* | GCTCAGTTTGGTGGTGTGGA | TCAAGTGGCTACGTTTGGTGT | 91 | 60 |
| *sars1* | TCAGCCAGTTTGACGAGGAG | CGATTGGCTGCTCTGATGTG | 111 | 60 |
| *cox8b* | GAGGGCTGCTATGAGACACC | TCACAAACATGACCGACAATGC | 113 | 60 |
| *atp5fa1* | GTCTGTCTGTGTCCCGTGTC | AGCAGCAACCTCACGATACT | 104 | 60 |
| *acsl5* | TGCTCTTTCACCATACCAGGAG | CACCATTGGATGCAAAGTAGTCC | 115 | 60 |
| *acot19* | TGAAACTAAACCCGGTCGCA | GAGAGGAGTGCGATGCTTGA | 115 | 60 |
| *fabp10a* | GCAGGTTTACGCTCAGGAGA | GCTGGATTTCTGTCACTGGC | 107 | 60 |
| **Primers for RNA probes** | | | | |
| | **F primer** | **R primer** | | |
| *atad3* - sense | AATTAACCCTCACTAAAGGG | TAATACGACTCACTATAGGG | | |
| *atad3* - antisense | GCGTGACACTCGTGTAGTGG | AGACTCCAGCACGGTCTGTC | | |
| **crRNA** | | | | |
| **crRNA target** | **crRNA sequence (PAM in bold)** | | | |
| *atad3* guide 1 | GCGCAGGAAAACCCTGAACG **AGG** | | | |
| *atad3* guide3 | ACGGTTCAGATGGAGCACCA **GGG** | | | |
